# MetaCherchant - an algorithm for analyzing genomic environment of antibiotic resistance gene in gut microbiota

**DOI:** 10.1101/106161

**Authors:** Evgenii I. Olekhnovich, Artem T. Vasilyev, Vladimir I. Ulyantsev, Alexander V. Tyakht

**Affiliations:** Federal Research and Clinical Centre of Physical and Chemical Medicine, Federal Medical and Biological Agency of Russia, Moscow, Russian Federation; ITMO University, Saint-Petersburg, Russian Federation; JetBrains Research, Russian Federation; Moscow Institute of Physics and Technology, Dolgoprudny, Russian Federation

## Abstract

Antibiotic resistance is an important global public health problem. Human gut human microbiota is an accumulator of resistance genes potentially providing them to pathogens. It is important to develop tools for identifying the mechanisms of how resistance is transmitted between gut microbial species and pathogens. We developed MetaCherchant - an algorithm for extracting the genomic environment of antibiotic resistance genes from metagenomic data in the form of a graph. The algorithm was validated on simulated datasets and applied to new "shotgun" metagenomes of gut microbiota from patients with *Helicobacter pylori* who underwent antibiotic therapy. Genomic context was reconstructed for several dominant resistance genes; taxonomic annotation of the context showed the species carrying the genes. Application of MetaCherchant in differential mode produced specific graph structures suggesting the evidence of possible resistance gene transmission within a mobile element that occurred as a result of the antibiotic therapy. MetaCherchant is a promising tool giving researchers an opportunity to get an insight into dynamics of resistance transmission in vivo based on metagenomic data.

## 1 Introduction

Spread of microbes resistance to antimicrobial drugs (antibiotic resistance, AR) is a global healthcare problem. Pathogenic microbes with multidrug resistance pose especially high hazard. According to the report of AMR [O*’*Neill 2016], the burden of AR-related deaths is predicted to increase to 10 million lives annually by 2050 and the global economical burden - to 100 trillion US dollars. The major factors contributing to the resistance spread is extensive medical and agricultural use of antibiotics [Rolain 2013].

Human gut microbiota is a reservoir of antibiotic resistance [Sommer 2009]. Under the influence of antibiotic consumption, gut resistome - composition of genes conferring AR from all microbes comprising microbiota [Wright 2007] - can increase in diversity and abundance (as assessed by quantitative metagenomics). Due to an active horizontal gene transfer within gut microbiota, the increase in resistome, amplified by the number of subjects in world population consuming antibiotics, strongly increases the chance for pathogenic microbes to obtain genetic resistance determinants from resistant commensal microbes inhabiting human body. Therefore, identification of resistome dynamics during and after antibiotic intake as well as the mechanisms of AR transmission within human gut is of utmost actuality.

Metagenomic analysis of gut resistome in populations of the world showed that national specifics of healthcare related to antibiotic usage and socioeconomic factors are reflected in the resistome composition as well as in the extent of its replenishment from the environment [Forslund; Pehrsson 2016]. Interestingly, significant levels of AR determinants were also detected in gut metagenomes of isolated populations having no access to antibiotics thus suggesting the global nature of AR transmission in microbial world.

Isolation and sequencing of individual bacterial genomes allows to examine genetic AR determinants of the strain in detail [Dai 2016] - particularly, to examine the genomic features surrounding the gene in order to assess the AR transmission history and potential. However, only a small fraction of microbes is cultivable, particularly, among gut-dwelling species. On the other hand, in the huge accumulated amount of human gut metagenomes, each dataset potentially contains information about all major species present in the community - thus making possible to approximately extract the data available from sequencing of an isolated strain. It can be performed at the general level of comparing relative abundance of AR genes [Yarygin 2016], as well as at a more detailed level - by exploring the genomic environment of an individual AR gene or operon. Common approaches to this task include metagenomic de novo assembly and subsequent analysis of contigs. The AR genes are identified in the contigs and their genomic context is analyzed to understand the location of the gene within a genome, the mobile element surrounding the gene and the surrounding of this island.

Such scenario works well in the case when the gene is present in a single species within a metagenome and occurs exactly once in a genomein single copy number per genomes. However, firstly, the genome might contain several AR gene copies. Secondly, gut microbiota is known to exhibit significant subspecies-level diversity (i.e. multiple subspecies of a single species with diverse genomes) [Greenblum 2015]. Thirdly, within a gut microbiota of a single subject, a gene can be present in several species simultaneously - which is likely to activate under the impact of antibiotics. The mentioned conditions suggest that during ordinary metagenomic assembly the linear contigs are likely to end at the location corresponding to genomic repeats and will provide only a ″planarized″ consensus image of the true genomic context of AR gene. Such poor representation does not allow to assess the environment correctly thus impeding the identification of the species - donor of AR gene - and the respective acceptor. From the perspective of personalized medicine, a more precise reconstruction of the AR evolution in vivo would improve the efficiency of resistance profiling for a patient and selection of optimal antibiotic therapy scheme. From the perspective of global healthcare, it would facilitate the tracking of significant trends in resistome spread as well as its control.

Here we present MetaCherchant - a novel method for exploratory analysis of genomic context of genes conferring antibiotic resistance directly from the metagenomic data based on local de Bruijn graph assembly. Unlike traditional assembly, MetaCherchant preserves the dimensional structure of this context, thus opening the way to a more accurate description of resistome dynamics and evolution in human microbiota. The method was validated using simulated datasets. Its application to gut metagenomes from patients before and after antibiotic therapy highlighted evidence of potential horizontal transfer of AR genes between different species.

## 2 Material and methods

### 2.1 Partial metagenomic assembly algorithm using a starting point

We developed a novel algorithm that performs classic steps of metagenomic assembly to the point of implicit construction of the de Bruijn graph and then builds a subgraph of the de Bruijn graph around a selected nucleotide sequence - the AR gene of interest. The algorithm allows to analyze the de Bruijn graph paths that contain the selected sequence thus making it possible to extract more information about the environment of the AR gene in the genome of one or multiple species within the microbiota. The algorithm was implemented basing on previously developed MetaFast software [Ulyantsev 2016] using Java programming language. The source code is available on GitHub (https://github.com/ctlab/metacherchant).

The first step of the algorithm is decomposing the input metagenomic reads into k-mers (nucleotide sequences of length k). The k-mers are stored in a hash table along with their coverage (the total number of times that particular k-mer has appeared in the reads). All the k-mers that appear with frequency below a fixed threshold are discarded as erroneous.

Due to the specificity of the problem the algorithm targets to solve, it is possible to overcome the classic k 31 restriction and, for larger values of k, only store the hash value instead of the actual k-mer. As the sequence of the target gene is known, it is only necessary to check if some specific k-mer was present in the reads without actually storing all the k-mers. This feature allows to increase memory efficiency of the algorithm, apply higher values of k thus providing high-detailed analysis of the graph structure with only moderate loss in performance. While this solution might produce undesirable hash collisions (because multiple k-mers can have the same hash value), the frequency of the latter is low compared to sequencing errors, and these collisions are easy to detect and fix.

Then a de Bruijn graph is constructed using the k-mers read from metagenomic data. Vertices in the graph correspond to k-mers and edges - to (k+1)-mers. Two vertices in the graph are connected by an edge if the nucleotide sequence on that edge is obtained by joining the vertices sequences overlapping by k-1. Genomic environment of a target gene is defined as some subgraph of the de Bruijn graph that contains that target gene.

To find that subgraph, we apply a modification of the standard breadth-first search (BFS) algorithm. In this algorithm, all visited vertices in the de Bruijn graph are stored in a queue and are processed in the order of extraction. It ensures that all vertices are added to the subgraph in increasing order of distance from the target gene. Therefore, the target environment subgraph is a set of k-mers closest to the target gene, and the sequences that are close to the target gene are likely to be close to the target gene in the metagenome itself. There are two possible stopping conditions: either the maximum amount of vertices in the subgraph is reached or the maximum distance from the target gene is reached, and the user can choose which one to use.

For convenient visualisation, a long non-branching path in a subgraph is displayed as one long sequence (also known as a unitig). It is achieved by the following algorithm: as long as there is a pair of vertices that can be merged, they are merged. Two vertices are merged when they are connected by an edge and have no other ingoing/outgoing edges. No edge in the graph is looked at twice, so the complexity is linear to the size of the graph. The resulting graph is saved in one of several formats including GFA (Graphical Fragment Assembly) and Velvet LastGraph. The graphs can be visualized with any program supporting those formats including Bandage [Wick 2015].

In single-metagenome mode (default), the algorithm processes a single metagenome to yield a single graph. A differential mode allows to compare the genomic environments of the same target genes between two different metagenomes - by constructing the combined graph from both datasets. When applied to paired datasets, this functionality allows the user to identify the changes in the environment, for example, differences in the environment of an AR gene in gut metagenome of a patient before and after antibiotics treatment (such changes indicate possible horizontal transfer event). The algorithm allows to find common and different parts of two subgraphs, detect overlapping subgraphs for different bacteria and postulate hypotheses about AR gene presence and transfer mechanisms.

**General workflow of environment construction algorithm**

~~~
**Input data**
[1] read metagenomic data and decompose them into k-mers.
**if** k ≤ 31 **then**
             save all k-mers in the hash table
**end if
if** k > 31 **then**
             save just hashes of k-mers in the hash table
**end if**
[2] find all k-mers included into the target gene sequence
**if** none of the target genes k-mers are present in reads **then**
             reported that it is impossible to construct the genomic environment
**end if**
[3] breadth-first search on the de Bruijn graph starting from k-mers of target gene
**if** either vertex limit in subgraph is reached or the graph radius limit is reached **then**
             stop
**end if**
[4] Compress non-branching paths of de Bruijn graph into unitigs
**Output:** Save the graph to file in one of supported formats
~~~

### 2.2 Data

#### Simulated data

For validation of the algorithm, simulated Illumina pair-end reads were randomly generated from selected microbial genomes using ART software [Huang 2012]. Single-genome (″genomic″) simulation was performed using *Klebsiella pneumoniae* HS11286 genome [Liu 2012]. Multi-genome (″metagenomic″) simulation was performed by randomly mixing the reads simulated from that and other four genomes: *Enterococcus faecium* EFE10021, *Escherichia coli* K12, *Bifidobacterium longum* BG7 and *Bacteroides υulgatus* ATCC8482 downloaded from NCBI GenBank database. In each of the simulations, targeted coverage for each of the genomes was 20x.

During the insertion simulations, the sequence of AR gene (or a mobile element including the gene) was inserted at a random location of a microbial genome. During the simulation of horizontal gene transfer event that occurred between first and second time points (corresponding to two metagenomes), the sequence of *K.pneumoniae* transposon Tn1331 was inserted into the genome of *K.pneumoniae* as well as of *E.coli.* The first metagenome was simulated from the transposon-carrying *K.pneumoniae* and transposon-free *E.coli,* and the second - from transposon-carrying *K.pneumoniae* and transposon-carrying *E.coli.*

#### Real sequencing data

The algorithm was applied to new ″shotgun″ metagenomes of stool samples collected from the patients with *Helicobacter pylori* before and after the *H.pylori* eradication therapy that included antibiotics intake [Gluschenko 2017]. The respective time points were denoted ″time point 1″ (before the therapy), ″time point 2″ (immediately after the therapy) and ″time point 3″ (1 month after the end of the therapy).

### 2.3 Data analysis and visualisation

Taxonomic profiling of metagenomes was performed using Metaphlan2 [Truong 2015]. AR genes were identified in the metagenomes by mapping the metagenomic reads to MEGARes database [Lakin 2016] using Bowtie2 [Langmead, Salzberg, 2012]. Relative abundance of the AR genes was calculated using ResistomeAnalyzer [Lakin 2016]. Taxonomic annotation of sequences corresponding to graph nodes was performed using Kraken [Wood 2014] and BLAST. Graphs were visualized in Bandage [Wick 2015]. Workflow of the data analysis described in the study is shown in Figure 1.

**Figure 1:**
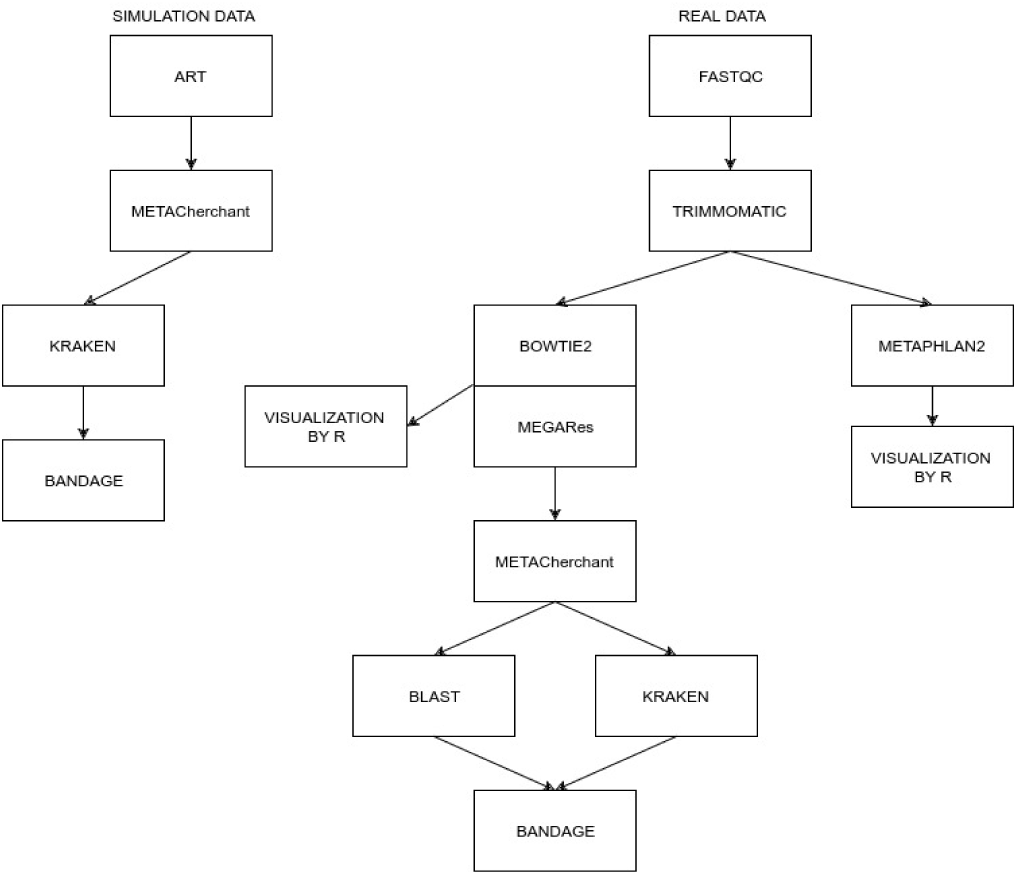
Workflow of the data analysis.

## 3 Results

### 3.1 Validation of the algorithm on simulated data

In order to check that the algorithm produces correct graph topology around the starting AR gene, we conducted a series of tests on simulated data of increasing complexity (see Figure 2).

**Figure 2:**
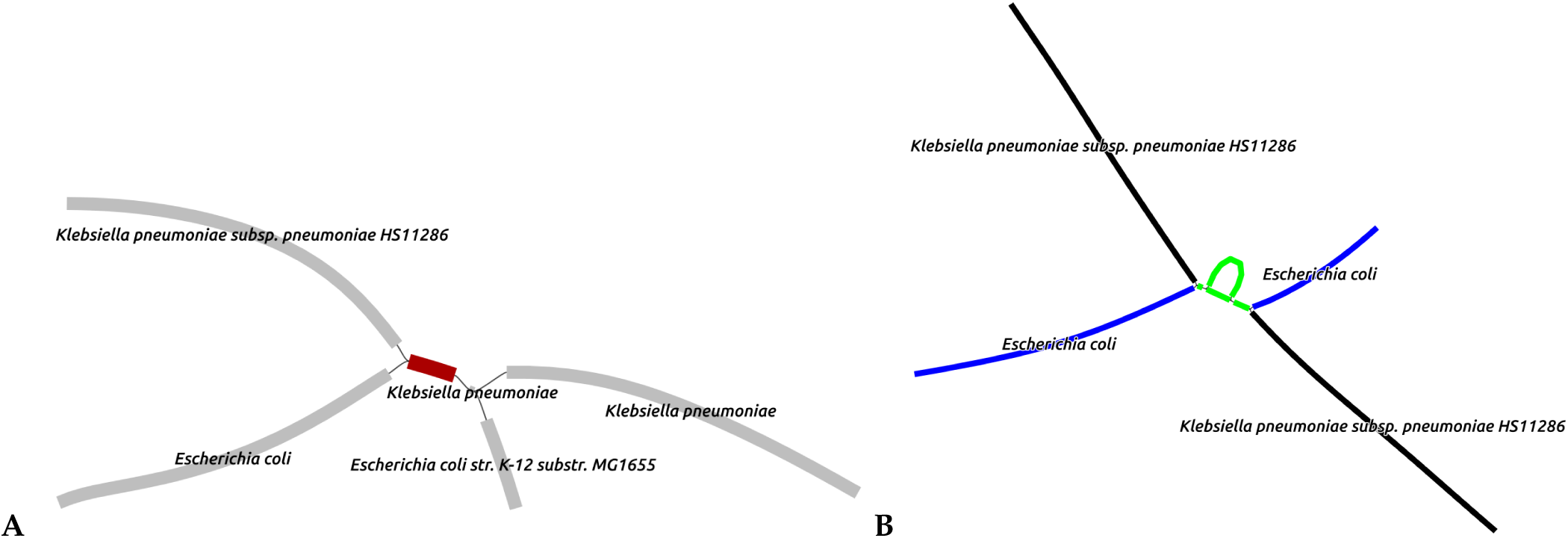
The output of algorithm produced from simulated sequencing data. B) Two-genome simulation (CTX-M gene in genomes *Klebsiella pneumoniae Escherichia coli*). C) Simulation of hypothetical horizontal transfer (the color of the fragment denotes: green - *Klebsiella pneumoniae* transposon Tn1331, black before and after antibiotics therapy, blue - only after).

During the first simulation, we reconstructed the genomic environment of a CTX-M (extended-spectrum beta-lactamase) gene naturally included in the genome of *Klebsiella pneumoniae* using simulated reads of this single genome. As expected, the produced graph was linear and the nodes flanking the target gene were annotated correctly. Secondly, in order to assess how the algorithm processes multiple occurrences of a gene in multiple species, the algorithm was applied to reads simulated from two genomes (*K.pneumoniae* and *E.coli*) with CTX-M gene sequence introduced into each of them. The obtained graph had a branching structure (Figure 2A); taxonomic annotation of nodes point at two unique ways of crossing the graph to assemble the fragments of the genomes containing the target gene. The results show that the algorithm correctly reconstructs the gene environment topology in the case when the gene is located in chromosomes of different species.

In the third simulation, we ran the algorithm in the differential mode to visualize a modeled horizontal gene transfer (HGT) of a mobile element containing AR gene from one bacterial species (*K.pneumoniae*) to another (*E.coli*) within gut microbiota (see Methods). The graph visualises the dynamics of genomic environment by combining the subgraphs from two time points before and after the HGT event and highlighting the differences between them (Figure 2B). Black color shows the sequences present in each of the metagenomes (they all belong to *K.pneumoniae*), while the blue color the sequences that were only in the second metagenome (they belong to *E.coli*). Thus, our algorithm allows to visualize the transmission of a mobile genomic element between the species. During subsequent simulations, the algorithm correctly reproduced gene environment for CTX-M gene and Tn1331 transposon for metagenomes generated from up to five bacterial genomes. The graph complexity increased with the number of genomes, and the values of k and minimum k-mer coverage were adjusted to achieve the most proper result.

### 3.2 Real gut metagenomes: analysis of taxonomic composition and resistome

Before analyzing the genomic environment of AR genes, we assessed the complexity of the gut metagenomes from the patients. As the result of taxonomic analysis, we detected 71 ± 18 species per metagenome in the analyzed gut metagenomes, signifying that the complexity of community structures is similar to one observed in human gut microbiota studies performed using similar approaches [Tyakht 2013]. also continued well represented many groups of AR genes were detected (53 ± 31 per metagenome). Basic results and types of bacteria resistance gene groups are shown in Fig. 3. These results suggest that the data are suitable for testing the algorithm.

**Figure 3:**
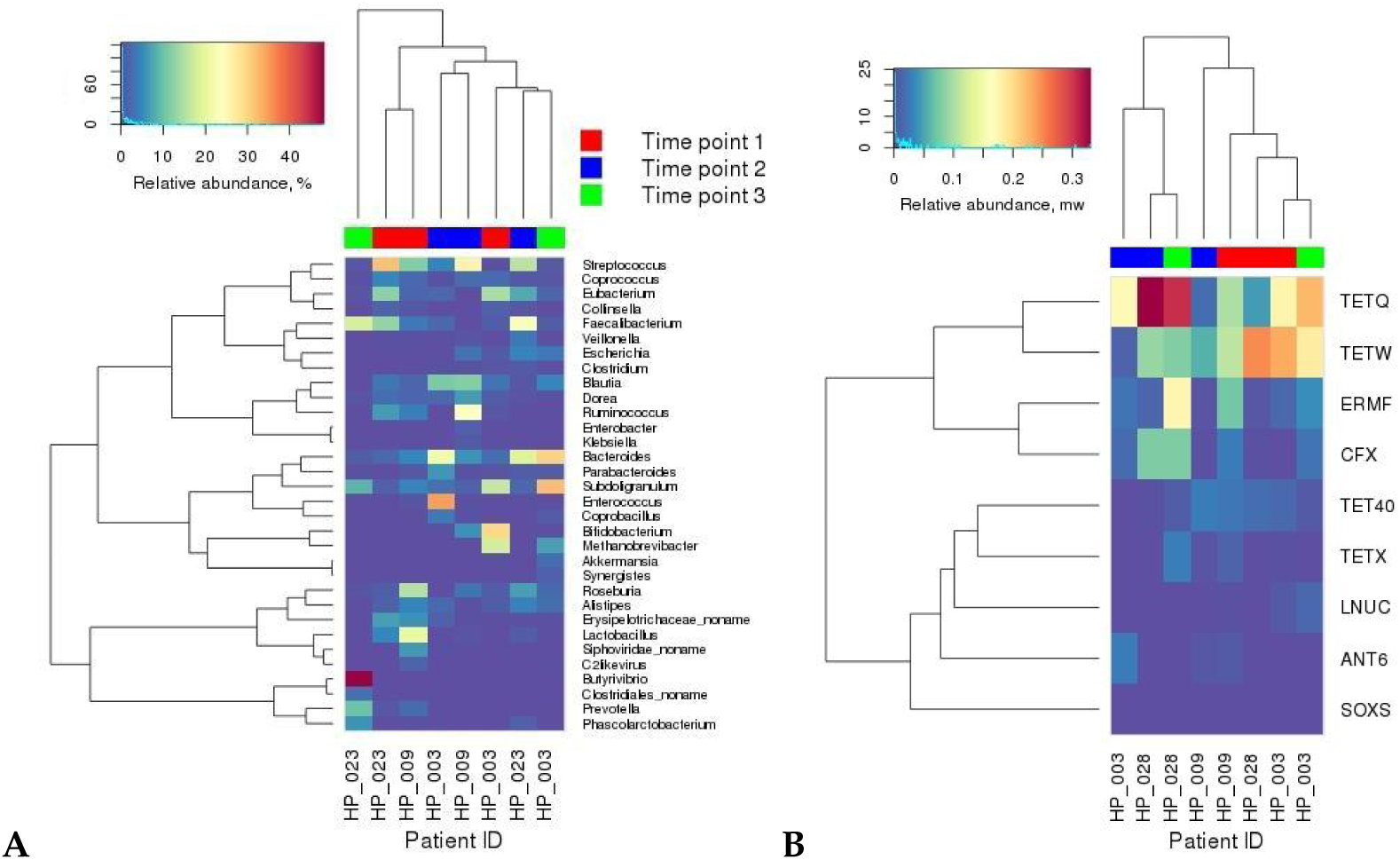
Relative abundance of the major bacterial species and AR gene groups in gut metagenomes. A) Relative abundance of major different bacteria species (percent) B) Relative abundance of major different group AR-genes (mass weight). The columns and rows were clustered using hclust function in R (Bray-Curtis distance (vegan package), ward linkage).

### 3.3 Real gut metagenomes: analysis of AR genes environment

MetaCherchant was used to explore the environment of the major AR genes detected in the metagenomic datasets. For each metagenome, genomic environment was constructed for each of the major detected AR gene groups mentioned above, using the most abundant gene per group (14 ± 8 genes per metagenome, totally 43 genes). We selected the most represented gene one per group. The analysis was performed over a range of control parameters (k = 41-141, minimum allowed k-mer coverage - 2-10×).

According to the results of our algorithm, some AR genes (or highly homologous genes) were detected in a genomic environment of a single bacterial species. An example is the adeC gene (multidrug resistance efflux pump) surrounded by sequences classified as *Enterococcus faecium* (Figure 4A). Some of the other genes were surrounded by environments from multiple species in the graph: the cfxA3 gene (class A beta-lactamase) together with short additional sequences were surrounded by two related but distinct species *Bacteroides vulgatus* and *Parabacteroides distasonis* (Figure 4B). We identified a structure homologous to transposon of *Streptococcus* spp. (*S.pyogenes* and *S.pneumoniae*) within the genome of *S.parasanguinis* that contained mel and msrD genes (macrolide resistance efflux pumps) (Figure 4C).

**Figure 4:**
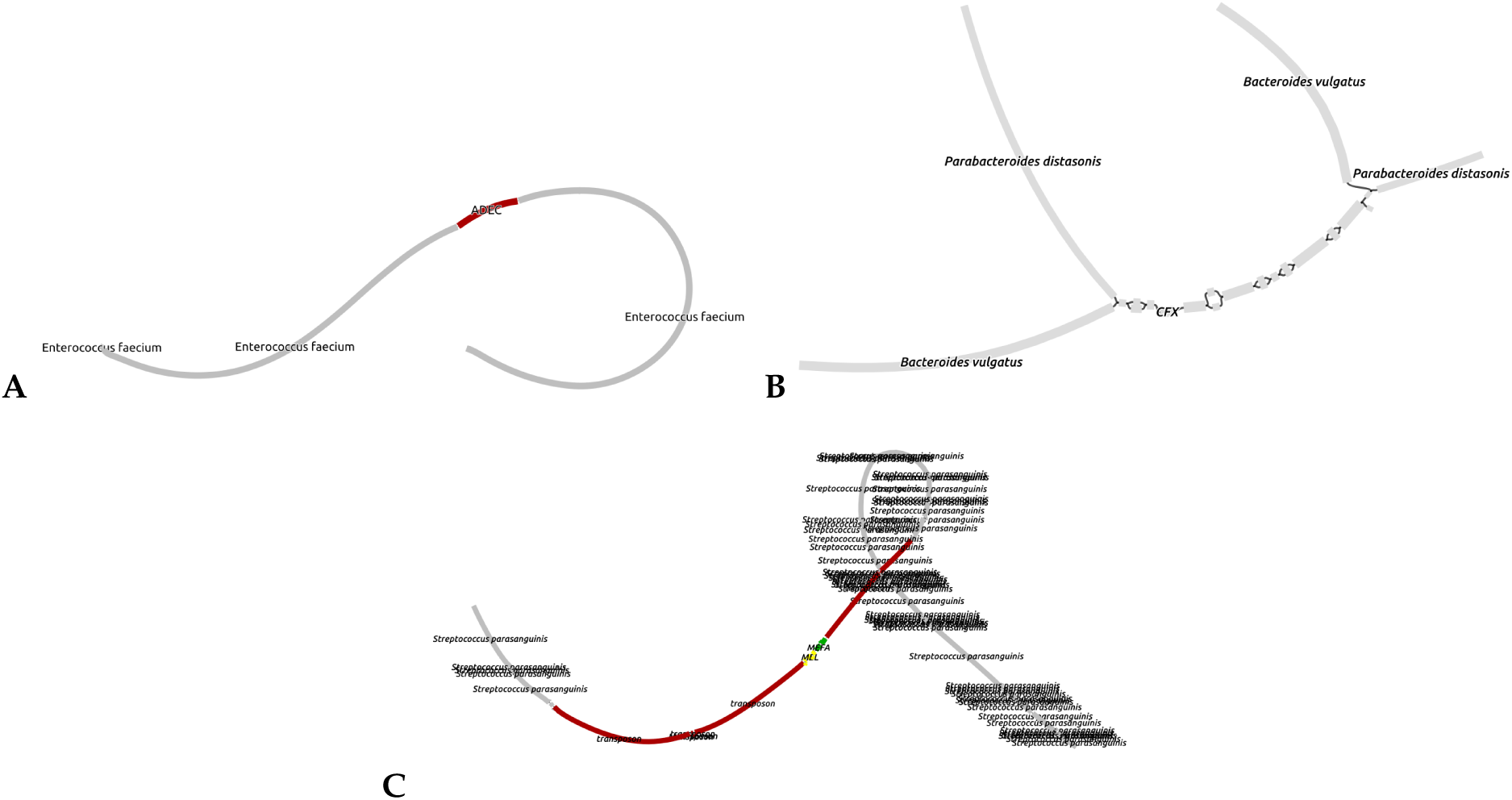
Genomic environment of AR genes reconstructed directly from real gut metagenomes. A) adeC gene in *Enterococcus faecium* genome B) cfxA3 gene in different intestinal bacteria genomes: *Bacteroides υulgatus* and *Parabacteroides distasonis* C) mel and msrD genes within trsnsposon-like structure in *Stereptococcus parasanguinis* genome

If an AR gene is a part of a large mobile element (longer than 70-100 Kbp), other mobile elements present in the microbiota are likely to contain sequences with high homology to that element. In such cases, topology of the obtained graph is complex and might include chimeric sequences and multiple adjacent resistance genes (an example is shown in Figure 5). Future implementations of the algorithm will allow to resolve the distribution of such genes among individual microbial species.

**Figure 5:**
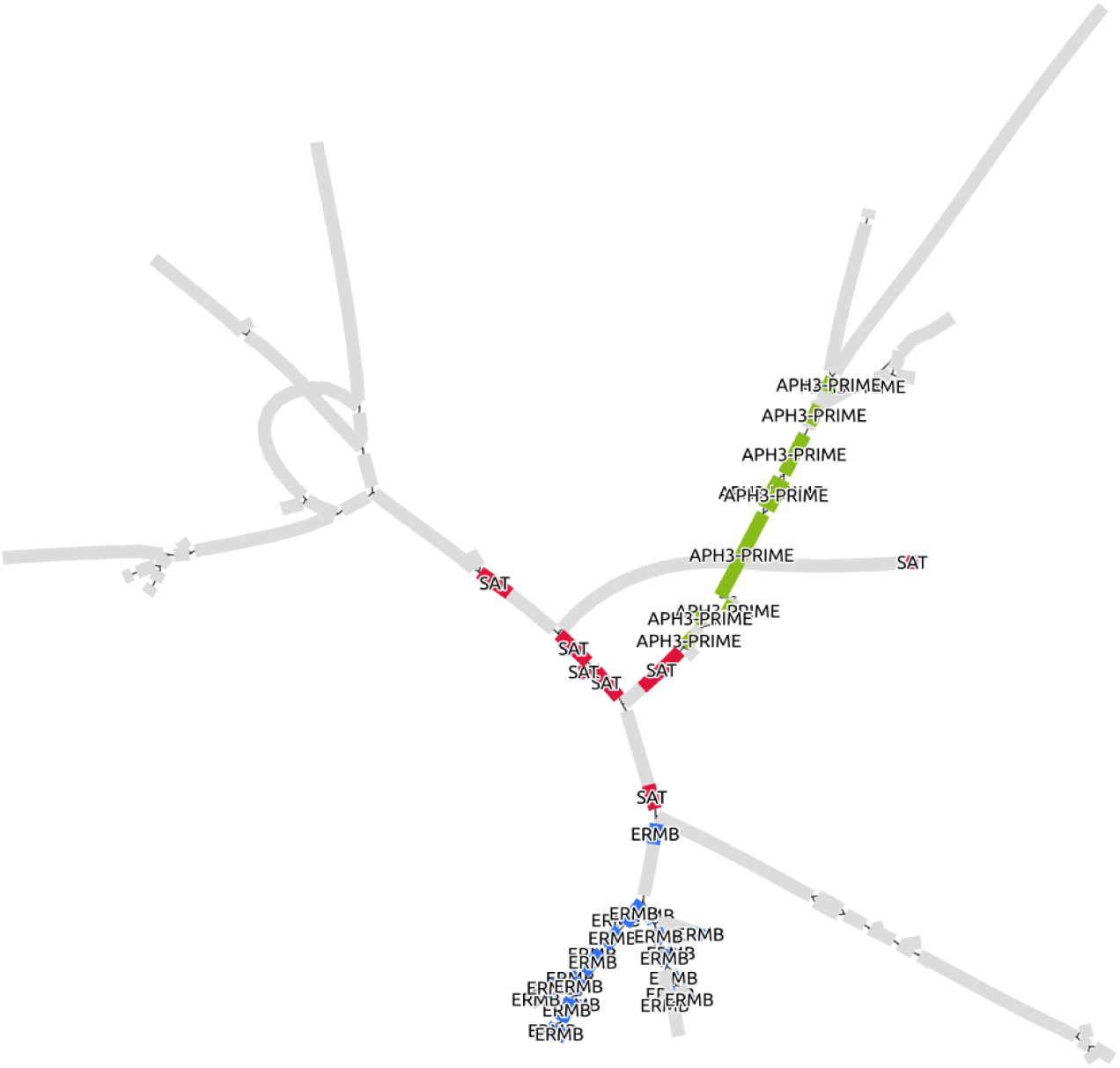
Reconstruction of genomic environment of APH3-PRIME gene showed co-localization of sat, ermB and APH3-prime genes in a plasmid-like context (sample HP009, time point).

### 3.4 Differential mode of MetaCherchant allows to identify possible events of AR gene transmission

Besides single-metagenome mode the examples of which are given above, it is possible to run MetaCherchant in differential mode allowing to overlay genomic environments of the same AR gene from several metagenomes. We applied differential mode to the paired gut metagenomes obtained from the same subjects before and after treatment to identify potential evidence of AR gene transmission. We processed the data from all metagenomes for each of the three subjects (HP003, HP009 and HP028, 14 ± 8 genes per metagenome). For some of the genes, it was possible to combine the environments across multiple time points.

An example of such case is displayed in Fig. 6: analysis of the gut metagenomes collected at two time points from a patient potentially shows a transition of ermF gene resistance from one genomic environment to another (red to blue, as shown in the figure), that is, appearing to be a HGT event. Taking into account the fact that the two analyzed time points correspond to the samples collected immediately after the H. pylori eradication course and one month afterwards, we speculate that the shown graph reflects the specifics of ″relaxation″ of the gut resistome following the end of antibiotic impact on gut microbiota.

**Figure 6:**
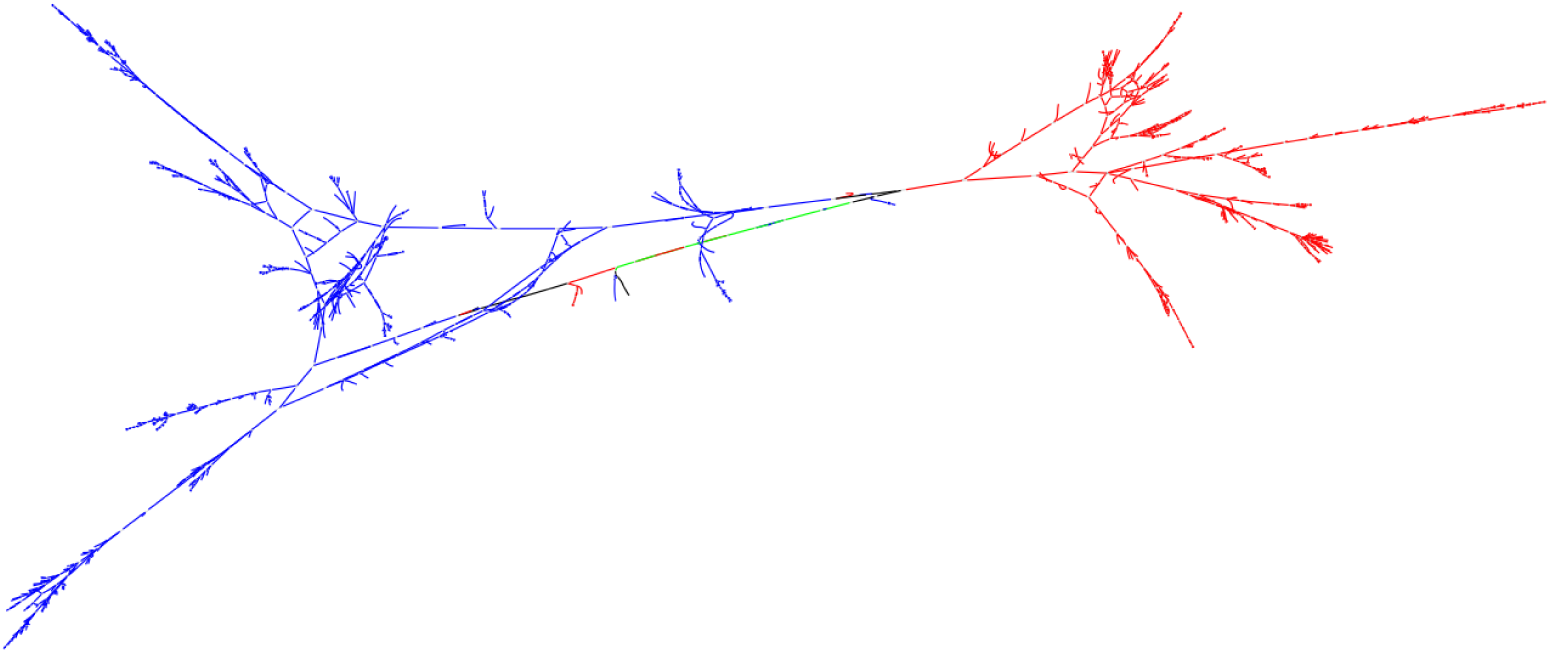
Combined graph of AR gene environment produced from two metagenomes of the same subject by running MetaCherchant in differential mode (patient HP028, ermF gene, time points 2 and 3). Red color denotes the part of the graph present only at the time point 1, blue color only at the point 2, black at both points; green color denotes the graph nodes corresponding to the target AR gene.

### 3.5 Performance

According to the described experiments, the total time for reconstructing a genomic environment of an AR gene in a single metagenome is about 15-20 minutes including download time (3-5 minutes), graph construction (3-5 minutes) and taxonomic annotation (5-10 minutes).

## 4 Discussion and conclusions

Gut resistome of healthy subjects is distributed among commensal bacteria and does not pose a threat by itself. However, in case of infection with antibiotic-sensitive pathogenic microbes, the AR genes could be transferred from normal microbiota, even from bacteria of a different genus, to the infectious agent. This phenomenon was observed in patients after antibiotic therapy [Karami 2007; Cremet 2012] and demonstrated experimentally on animals and healthy donors [Lester 2004; Lester 2006]. There were events of hospital infection expansion, causative agent of which obtained AR from patients gut microbiota [Goren 2010; Cremet 2012]. Thereby, gut microbiota represents an important reservoir of AR genes open to infectious agents of socially significant diseases.

The proposed algorithm allows the user to explore the connections between AR genes and genomes of taxa present in microbiota using a graph representation. Unlike the existing methods, MetaCherchant provides a rich representation of genomic context of AR gene, thus showing the resistance potential of species in gut microbiota in an unbiased way, as well as providing means for examining the ways of transmission of resistance. The method can be applied not only to gut metagenomes, but also to other environments it is especially important in the light of discovered transmission of resistance to gut from urban environment [Pehrsson 2016]. Certain limitations of the method are linked to the complexity of the graph structures arising when the AR gene is within a mobile element prevalent in a high number of species. It is sometimes difficult to accurately determine the AR gene recipient and donor species because plasmids from different species may contain highly homologous conserved regions leading to the formation of chimeric structures. Our method will contribute to the design of rational antibiotic therapy schemes for infectious diseases treatment (including the sequence of use of known drugs and introduction of new antimicrobial drugs [Imamovic 2013]). This will provide both increase of success rate for individual patients and constrain the spread of new multidrug-resistant pathogens.

## 5 Acknowledgements

Algorithm testing and biological application were financially supported by the Russian Scientific Foundation (grant 15-14-00066). Algorithm development was financially supported by the Government of Russian Federation (grant 074-U01).

